# Nano-hydroxyapatite coating synthesized on quasi-fibrillar superstructures of collagen hydrolysate leads to superior osteoblast proliferation when compared to nano-hydroxyapatite synthesized on collagen fibrils

**DOI:** 10.1101/2021.02.13.431097

**Authors:** Pradipta Banerjee, Mayur Bajaj, Chetna Bhat, Y Geethika, Hemanth Irle

## Abstract

This study had a two-fold objective: To utilize collagen hydrolysate for synthesizing a nanoscale Hydroxyapatite (HA) coating that would act as a superior osteoblast adhesion/proliferation agent compared to collagen-derived HA (C/HA) and to comprehend the significant role played by structural constraints on HA nucleation. Collagen was extracted from pacu skin with a high yield of 65.3% (w/w of tissue). It was digested by collagenase and the hydrolysate (CH) was purified with a high yield of 0.68g/g of collagen. The CH peptides had a mass of 6kDa, a predominant PP-II conformation and formed self-assembling hierarchical structures at physiological pH with dimensions of 842.2±229nm. The HA synthesized on CH (CH/HA) displayed higher yield when compared to C/HA. Structural analysis of CH/HA revealed that the PP-II peptides coiled to form mimic-helical moieties with reduced intermolecular packing distance of 0.9nm. The mimic helices cross-linked to form a vast quasi-fibrillar network that was comparatively smaller than collagen fibrils but exhibited enhanced stability and greater dynamicity. CH/HA displayed intense calcium-carboxyl interactions, sharper diffraction planes, smaller size of 48±6.2nm and a Ca/P ratio closer to 1.69 when compared to C/HA along with displaying serrated edge blooming crystals. Because of the small size, the CH/HA nanocrystals displayed significantly better osteoblast adhesion than C/HA and reduced the doubling time of cells. Overall, the results indicated that CH based nanocomposites displayed suitable morphological characteristics and cellular response for potential application as implant and bone graft coating material.

**Graphical abstract:** 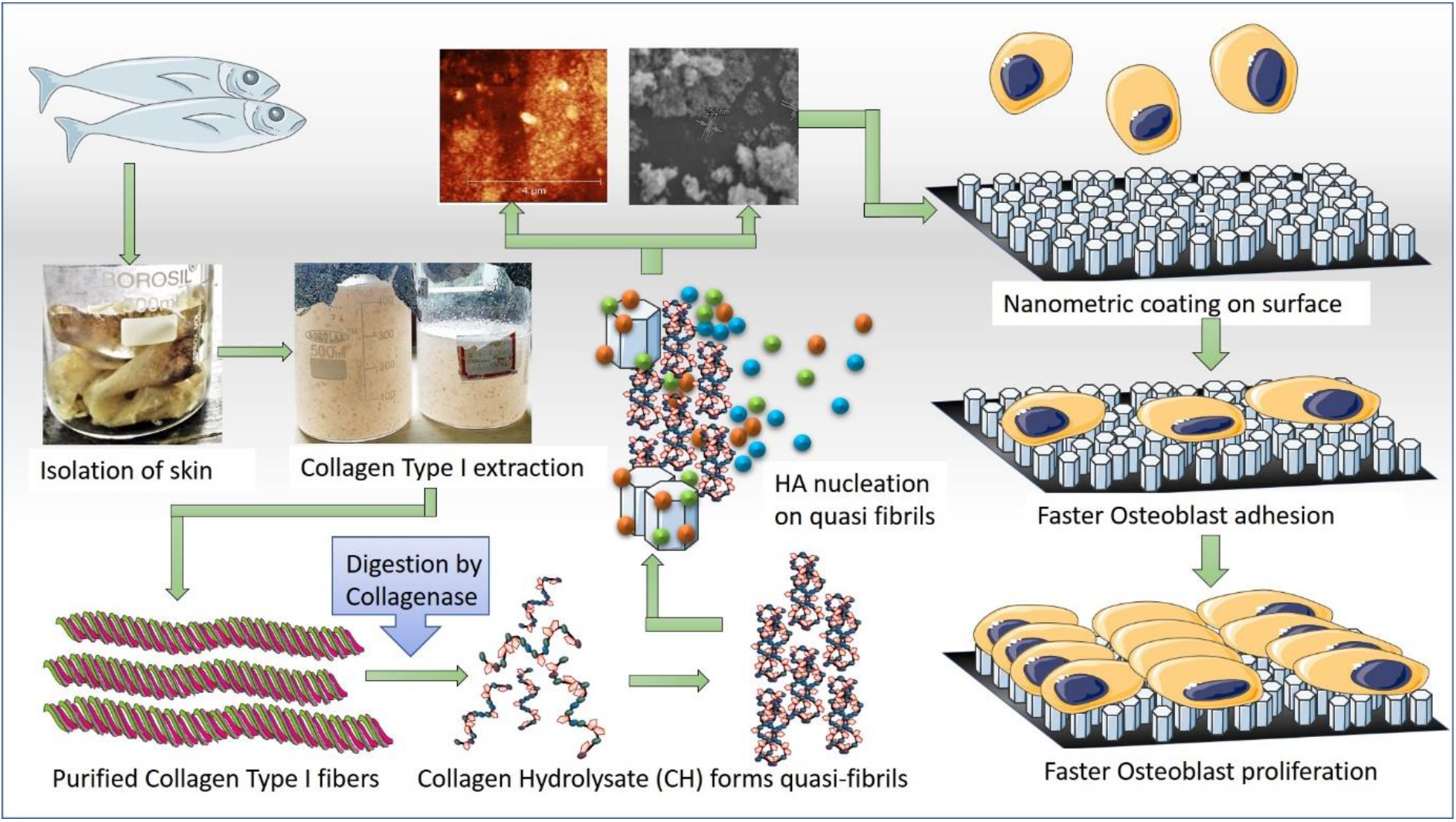

## 1. Introduction

Orthopedic implants are an effective treatment to repair fractures and replace lost portions of bone tissue. [1]. However, a substantial number of implants fail to integrate with host bone owing to their inability to activate osteoblast proliferation, leading to a considerable number of rejections and revision surgeries [2,3]. A customary technique of constructing host-friendly endosseous metal components involves coating the surface with a bioactive and osteoconductive material like hydroxyapatite (HA) [Ca_10_(PO_4_)_6_(OH)_2_] [2,4]. However, chemically synthesized HA displays deformed crystals, low tensile strength, and a variable composition depending on its scheme of synthesis, consequently leading to incoherent comparing parameters [5,6].

Biomimetic strategies offer a simple and effective solution to create HA [7]. The procedure requires suspension of a protein substratum, generally collagen type-I, in a HA nucleating solution allowing *in vitro* crystal formation. The reaction process is easy to control and offers a high yield. Even though the method has been known for over a decade, the exact mechanism of *in vitro* HA nucleation is yet to be deciphered in its entirety [8]. Knowing the mechanism is imperative as it will allow high-throughput synthesis of HA with dimensions similar to bone crystals, which, when coated on medical implants, will guide faster osteoblast adhesion, leading to effective osseointegration.

The current theories that attempt to decipher biomineralization can be divided roughly into two doctrines. One school of thought pursues that the presence of negatively charged groups is sufficient for biomineralization to occur. In this theory, the presence of acidic residues or carboxyl groups is sufficient to nucleate HA, superseding the need for collagen or any structural hierarchy [9,10,11]. The studies that champion this thought process also suggest that in physiological conditions, nucleation is initiated by acidic residues from accessory proteins while collagen merely supplies the structural framework necessary for crystals to propagate [12]. The alternate theory considers the structural aspect of collagen Type-I to be of primary importance. It suggests that the triple-helical nature of collagen and its long fibrillar arrangement initiates and regulates the interaction between terminal amino acid residues and inorganic ions resulting in HA formation [13]. In this theory, the process of mineralization is heavily influenced by the secondary structure with amino acid sequence playing a comparatively less significant role [14,15,16].

It was clear from a thorough literature survey that an attempt by either process would lead to one result or the other but not a definitive solution. So, the authors tried a different approach by *attempting to nucleate HA on collagen hydrolysate (CH) instead of collagen*. As of today, there exist only two studies that have mentioned HA nucleation on CH. One among them lists the conditions that would allow CH from chrome tanning waste to nucleate HA [17]. However, the yield, dimensions, and osteophilicity of the crystals have not been reported. The other study reports the use of CH, but only as an accessory source of carboxylic acid in the biomineralization of collagen [18].

Here, we report the first study on the bio-mimetic mineralization efficacy of fish collagen hydrolysate in the absence of any accessory proteins. We show that the constituent peptides of collagen hydrolysate are self-sufficient in initiating and orienting the formation of carbonated HA *in vitro* in a manner that is similar to bone formation in the body. We also report that CH is significantly more efficient than collagen in producing HA crystals that possess sharper crystal planes and are smaller in dimensions leading to higher osteoblast adhesion and proliferation. The results obtained in this study provides substantial insight into the mechanism of HA nucleation and help in understanding the conditions governing *in vitro* biomineralization. Consequently, this leads to a deeper understanding of the *in vivo* HA nucleation process, opening up pathways to synthesize protein-HA composites with smaller dimensions and higher osseointegration properties to be used as coating on medical implants.

## 2. Materials and Methods

The collagen used for this study was collected from fish processing by-products. The chemicals utilized in the experiments were procured from Himedia Labs Pvt Ltd., Mumbai. Collagenase type-I enzyme was purchased from Juniper life sciences, Bangalore. Antimycotic/Antibiotic solution and Glutaraldehyde grade 1 were procured from Sigma Aldrich.

### 2.1 Isolation of collagen

The fish species to be used for collagen extraction was chosen based on a market survey along with an analysis of FAO reports. Red-bellied Pacu is one of the fastest-growing freshwater fish species released in India [19,20]. During the time of the study, it was sold in high demand in local markets and hence its processing by-products were chosen for the source of collagen extraction. Descaled skin samples were collected from a nearby local market, brought to the University, washed and dry weight of the material was noted. Acid soluble collagen was extracted following a standard protocol [21]. The extraction involved successive treatments by 5% NaOH, 5% butanol, and 0.25% EDTA to remove non-collagenous proteins, fats, and minerals respectively with dry weight being noted between every successive treatment. The post-treatment material was dissolved in 0.5M acetic acid for 36 hours with regular changes of the solution. This cycle was conducted repeatedly to extract the maximum amount of collagen. The extracts were pooled and collagen was precipitated by adding 6% (w/v) NaCl. The precipitated collagen was desalted by dialysis against deionized water for 72hrs and dried in a VD-250R Freeze dryer (Taitec, Japan).

### 2.2 Hydrolysis of collagen and purification of CH

The lyophilized collagen sample was subjected to fragmentation using bacterial collagenase. Collagenase type-I was dissolved in 0.1M phosphate buffer, pH 7.2, at a concentration of 4mg ml^−1^. This was added to a solution of collagen (100mg ml^−1^) dissolved in an identical buffer system. The reaction mixture was incubated at 37°C for 24hrs. The solution was centrifuged at 5000rpm at 6°C for 15min to remove any suspended debris. The supernatant was mixed with acetate buffer, pH 4.5, and applied to a 1.5 × 5cm CM Sepharose column attached to a low-pressure automated chromatography unit. The sample was eluted with a linear gradient of NaCl ranging from 0 to 1M. Fractions representing the eluted samples were pooled and applied to a 1.5 × 3cm PD-10 column. Deionized water was run, and desalted CH sample was collected. The pooled sample was freeze-dried to obtain a powder.

### 2.3 Structural attributes of CH

#### 2.3.1 SDS-PAGE of CH

Electrophoresis was carried out according to the method of Laemmli [22]. CH (100μg ml^−1^) was run in an 8% resolving gels. Intact skin collagen, rat-tail tendon collagen, and known molecular weight markers were run under similar conditions for reference. The gels were stained as per standard protocols [23].

#### 2.3.2 Folding studies on CH

The stability of CH was checked by measuring the pH-induced shift of the 230nm peak. The samples were dissolved at a concentration of 10μg ml^−1^ in 0.1M of numerous buffer systems, including acetate (pH 4 and 5), citrate (pH 6), phosphate (pH 7 and 8), and tris buffer (pH 9). Absorbance spectra were analyzed by scanning in the range of 200-400nm using a Genesys 10S UV-Vis Spectrophotometer (Thermo). Collagen was used as control.

#### 2.3.3 Zeta potential and dimensions of CH

A Nanotrac wave II Q was used to measure the zeta potential and dimensions of CH in 0.1M phosphate-buffered saline, pH 7.4. Background measurement was performed using only buffer. Collagen was used as control.

#### 2.3.4 Circular Dichroism studies on CH

CD studies were conducted to determine the effect of pH on the secondary structure. CH was dissolved in 0.1M phosphate buffer, pH 7.4 at a concentration of 2mg ml^−1^. The solutions were centrifuged at 8000rpm for 1min to obtain a clear supernatant. 400μl of the supernatant was loaded into a quartz cylindrical cuvette with a path length of 1mm. A Jasco J-715 spectropolarimeter was used to carry out the spectral analysis. Collagen, treated similarly, was used as control.

#### 2.4 Biomimetic growth of HA crystals on CH

HA crystals were bio-mimetically synthesized on CH by following the protocol of Banerjee et al. [17]. CH (500 mg) was suspended in a solution containing 100μmol ml^−1^ of NaCl and 10μmol ml^−1^ of Na_2_HPO_4_. 20 sets were made. Each set was sealed after adding 150μl of 1:10 diluted antimycotic solution and 2.8μl of glutaraldehyde grade 1 (25% in H_2_O). After 24 hours, 20μmol of CaCl_2_ dissolved in phosphate buffer of pH 7.4 was added to all the sets. The tubes were re-sealed and incubated at 37°C for 31 days for crystal growth. The crystals were washed 6 times in distilled water and freeze-dried as before. The dried crystals were subjected to structural analysis. Crystals grown on collagen, treated similarly, were used as control samples. The composites were labelled as CH/HA and C/HA for HA synthesized on CH and collagen respectively.

### 2.5 Characterisation of CH/HA crystals

#### 2.5.1 Photoluminescence profile of CH/HA

Fluorescence intensities of CH/HA were measured by a Cary Eclipse Fluorescence Spectrophotometer. The emission spectrum was detected at a voltage of 800V, in the range of 240-550nm with a scanning rate of 300nm min^−1^.

#### 2.5.2 IR Spectral profile of CH/HA

The functional groups of the crystals were profiled using Thermo Nicolet iS5 FT-IR. 200mg of KBr was ground with 1mg of CH/HA crystals to form a translucent pellet. The pellet was utilized to read the transmittance profile at the IR range.

#### 2.5.3 XRD profile of CH/HA

XRD analysis of CH/HA was performed on Rigaku D/Max X-ray diffractometer with Cu Kα radiation source (wavelength = 1.54Å). The supplied voltage and current were set to 40kV and 120mA, respectively. The samples were exposed at a scan rate of 10° min^−1^ from 5° to 60°.

#### 2.5.4 Zeta potential and dimensions of CH/HA

A Nanotrac wave II Q was used to measure the zeta potential and size of CH/HA samples. Background measurement was performed using water. The sample was suspended in deionized water and measurements were taken for 30sec. C/HA was used as control.

#### 2.5.5 SEM-EDX analysis of CH/HA

SEM was performed on CH/HA crystals to investigate their morphology and porosity. CH/HA crystals were sputter-coated with gold and were observed under a field-emission scanning-electron microscope with a magnification of 1500X (S-4700, Hitachi Ltd., Tokyo, Japan). The interfaces were examined in secondary electron mode at 1keV accelerating voltage and 10mA emission current. SEM images were acquired digitally by a data acquisition system. Energy-dispersive X-ray spectroscopy (EDX) was used at specific locations to detect the Calcium/Phosphate ratio (Ca:P) and their distribution within the sample.

#### 2.5.6 Dimensional analysis of CH/HA

The topological assessment and crystal size estimation was determined by atomic force microscopy (AFM). The crystals were dispersed in 50% ethanol and dispensed on a mica coverslip. The coverslip was placed inside a desiccator and left undisturbed until it dried. AFM analysis in Non-Contact Mode was conducted for the dried sample using an A-100 AFM (APE Research, Italy).

### 2.6 Osteoblast adhesion/proliferation on CH/HA

The osteosarcoma cell lines (MG-63) were cultivated according to the recommendation of the supplier, in DMEM containing 10% FBS and 40 IU ml^−1^ of antibacterial and antimycotic solution. Cells were cultured in T25 flasks in a humidified 5% CO_2_ incubator at 37°C. For passaging, cells were detached with trypsin/EDTA and subsequently replated. Trypan blue exclusion assay was carried to check the viability of the cells and cell count was estimated using Neuber’s chamber.

The test (CH/HA) and control (C, CH, C/HA) samples, in triplicates, were coated on 48 well plates at an amount ranging from 10μg to 1mg per well. The plates were exposed to UV for 3 hours, followed by air-drying, rinsing with 0.1M PBS and re-drying. MG63 cells suspended in DMEM media were seeded on to the test and control wells at a count of 4×10^4^ cells per well. The plates were kept in a CO_2_ incubator for 6 hrs and cell adhesion measured by MTT assay.

Cell proliferation assay was conducted in triplicates. A fixed number of cells were added to the coated dishes and an MTT assay was conducted every 12h for 72h. A calibration curve was prepared with a known cell count. The doubling time of the cells grown was calculated by fitting the curves to the exponential function given below by the least-squares method.

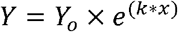

where Y and Y_0_ signify the change in cell count with time and the cell count at time zero respectively, x denotes the time in hours and k is the rate constant to be determined, expressed in h^−1^. The doubling time (t) was calculated by the equation:

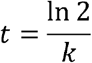

The datasets were analyzed by two-way ANOVA with post hoc Tukey’s Test. P values less than 0.05 were considered significant.

## 3. Results and discussion

### 3.1 Isolation and Extraction of collagen

Table 1 depicts the % of acid-soluble collagen (ASC) extracted from the skin of *Piaractus brachypomus* (Red-Bellied Pacu). The extraction process yielded 65.3% of collagen. The results matched with collagen yield obtained from Catla and Rohu skin (65-69%) [24] and was higher than most acid extraction techniques [25–29].

**Table 1 -.**
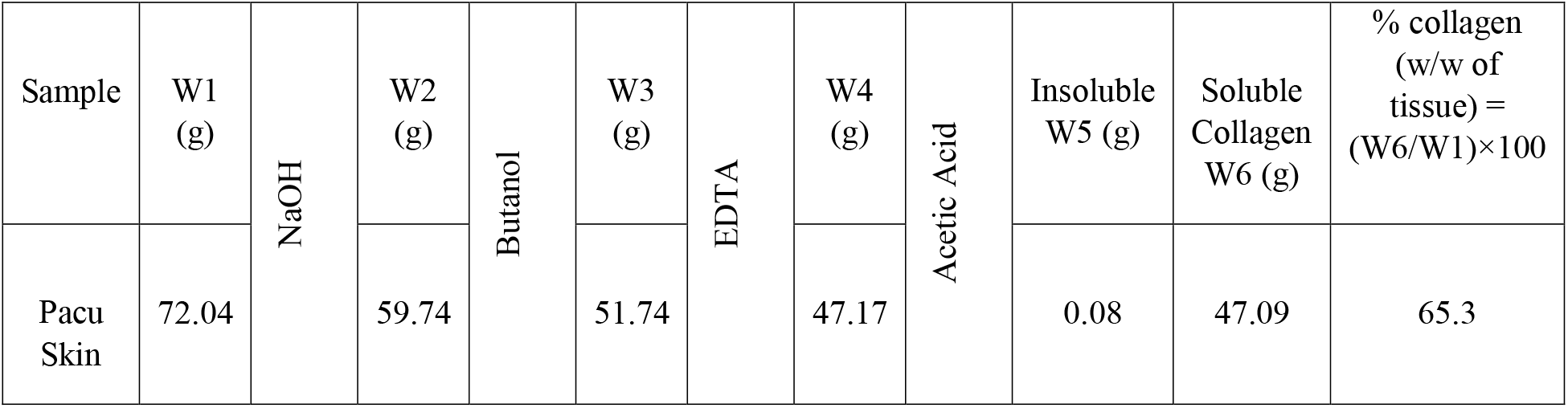
Isolation of Collagen type 1 from Pacu skin. The dry weights in grams were noted as W1, W2, W3 and W4 for skin, after cleaning, NaOH, Butanol and alkaline EDTA treatment respectively. W5g accounts for insoluble material after acetic acid treatment while W6g represents soluble collagen.

| Sample | W1<br>(g) | NaOH | W2<br>(g) | Butanol | W3<br>(g) | EDTA | W4<br>(g) | Acetic Acid | Insoluble<br>W5 (g) | Soluble<br>Collagen<br>W6 (g) | % collagen<br>(w/w of<br>tissue) =<br>(W6/W1)×100 |
| --- | --- | --- | --- | --- | --- | --- | --- | --- | --- | --- | --- |
| Pacu<br>Skin | 72.04 |  | 59.74 |  | 51.74 |  | 47.17 |  | 0.08 | 47.09 | 65.3 |

The efficiency of collagen extraction depends heavily on the extraction protocol and to some extent, on fish species, source tissue, and environmental conditions [30]. A large number of studies use pepsin along with acetic acid to increase the efficiency of extraction [31,32]. Pepsin cleaves the cross-linked telopeptides of collagen leading to better solubility and higher yield [30]. However, enzymatic treatment is costly and leads to bottlenecks in large scale extraction. At the same time, pepsin-mediated removal of telopeptides and associated sequences could lead to a loss of potential HA nucleating sequences. The higher yield obtained in this study, even without the use of pepsin, was due to specific modifications of the extraction protocol that included the use of milder NaOH during pre-treatment and repeated acetic acid extraction. Multiple cycles of ASC may appear as time-consuming, but it ensures greater extraction of intact collagen at a lower cost.

### 3.2 Hydrolysis of Collagen type-I and Purification of CH

The complex hierarchy, steric crowding, and ample abundance of proline in collagen renders it insusceptible to complete degradation by digestive protease. Efficient degradation can only be possible through bacterial collagenolytic enzymes which cleave at specific repetitive sites [33]. The isolated collagen was fragmented by Collagenase type I-A from *Clostridium histolyticum*, which cleaves at the repeating sequence P-X-G-P where X is a neutral amino acid.

The CH exhibited an elution profile as shown in Fig. 1a. A prominent peak was observed at 43.50% NaCl for CH. The higher ionic strength required for elution reflected the higher charge density of CH due to the increased number of free N and C-terminals. Contrastingly, the control collagen sample eluted at 8.2% and 28% NaCl. The latter peak was attributed to the elution of α chains whereas the former small peak was attributed to higher-order structures formed by the recoiled α-chains, a signature profile of collagen type-I. The CH yield was calculated to be 0.68 g/g of collagen, which was comparatively higher than previous reports [27].

**Fig. 1.**
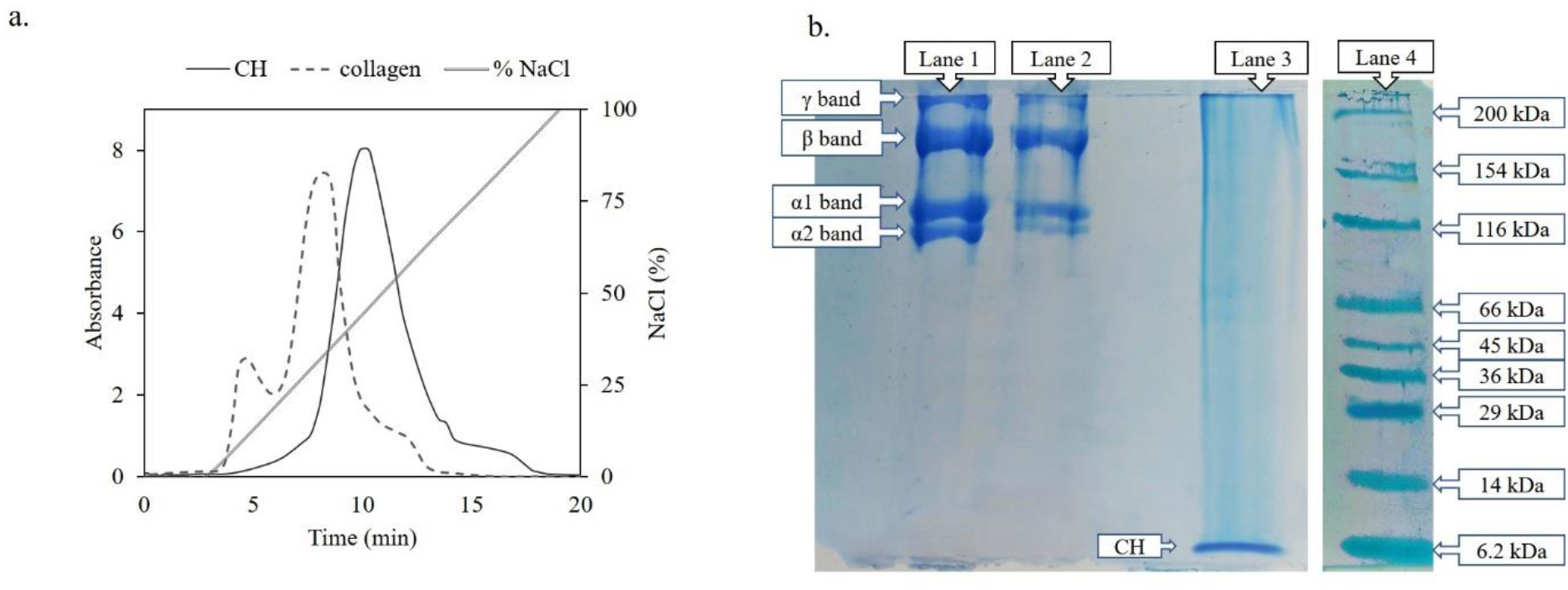
**a.** Purification of CH from collagen type 1 by IEC. A shoulder peak at 8.30% and an intense peak at 43.50% of NaCl concentration indicating the presence of aggregated collagen and purified CH respectively. **b.** Electrophoretic profile of isolated collagen (lane I), rat-tail tendon collagen (lane 2, control), collagen hydrolysate (lane 3) and standard marker (lane 4).

### 3.3 Molecular weight profiling of CH

As shown in Fig. 1b, the electrophoretic pattern exhibited by the isolated collagen (Lane 1) matched with that of rat-tail tendon collagen (Lane 2), confirming it to be of type-I [34]. The unique recoiling nature of collagen is depicted in its signature electrophoretic profile with two prominent α bands, resulting from its twin α1 chains and single α2 chain, along with higher-order β and γ band [35]. The molecular weight (MW) of α1 and α2 bands corresponded to values of 125kDa and 110kDa respectively. The presence of the higher-order β and γ bands are attributed to recoiling of α chains with each other, a pattern also observed in the ion-exchange profile of collagen. Lane 3 depicted the enzymatic digest of collagen run under similar conditions. The CH was devoid of any α or higher-order profiles and displayed a low molecular weight band at the bottom of the gel. The MW, when compared to standard marker run under similar conditions (Lane 4), was determined to be around 6kDa with the possible presence of fractions with even lower MW.

### 3.4 Structural attributes of CH

An intense peak at 220-230nm corresponds to the presence of amine groups in peptide bonds. This peak has been known to corroborate the structural state of the protein and indicates hierarchical assembly formation [36]. A hypsochromic shift of this peak with changing pH provides evidence of folding characteristics of biomolecules.

Fig. 2a exhibits the pH-induced spectral shift of CH. The profile can be compared with the folding tendencies of intact collagen type-I as shown in Fig. 2b. At pH 4, CH exhibited an absorbance maximum (or λ_max_) at 230nm while collagen displayed a broad peak at 221nm. The difference in λ_max_ was attributed to the absence of folded states in CH [37]. Interestingly, upon changing the pH from 4 to 5, CH underwent a blue shift indicating that its peptide components were able to orient themselves in a hierarchical proto-aggregated structure. Collagen retained its absorbance maxima and correspondingly, its structural state.

**Fig. 2.**
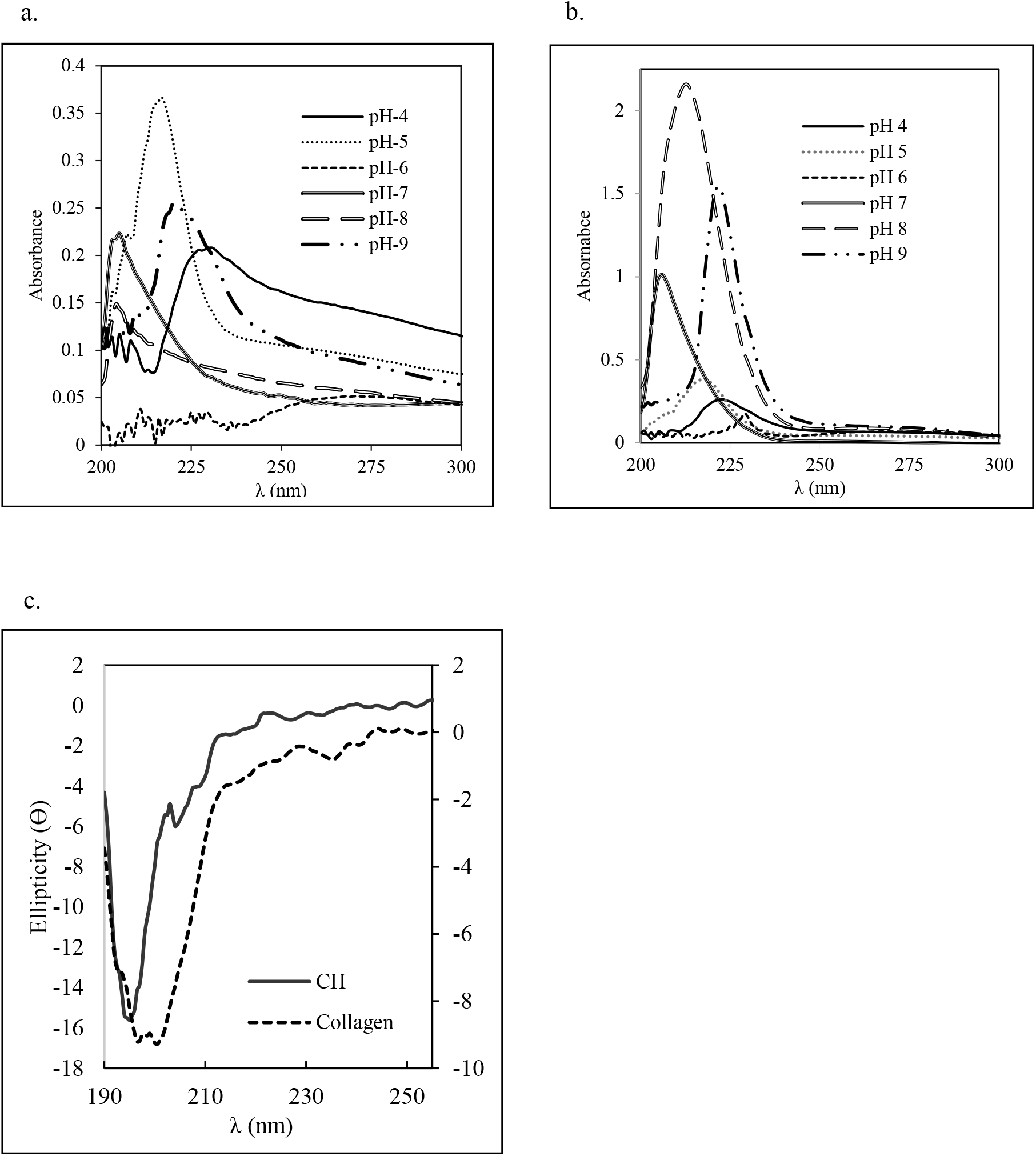
**a** – pH-dependent Absorbance profile shift of CH displaying tendency to self-assemble at neutral and alkaline pH **b** – pH-dependent absorbance profile shift of collagen displaying fibril-formation tendency at pH 7 **c** - CD spectra of CH and collagen at pH 7.4 with the former exhibiting a tendency to form fibrils

At pH 6, both CH and collagen displayed low solubility and a redshift at 230nm. This was due to the fact that the α1 chain of collagen has a pI nearing 6, rendering it with zero net charge and low solubility. This results in distortion of the proto-aggregated structures leading to the uncoiling of the components. Also, presence of the insoluble protein chain leads to scattering of incident wavelengths decreasing the signal-to-noise ratio of the spectrum.

A sharp blue shift was observed at pH 7 for both CH and collagen resulting in a λ_max_ at 205nm. The shift demonstrated the tendencies for both CH and collagen to fold into higher-order structures. Collagen is known to arrange into a fibrillar assembly at physiological pH, explaining its blue shift [37]. It was possible that the peptides in CH oriented themselves to form a sort of quasi-fibrillar structure, mimicking the fibrillar structure of collagen. As the pH was made more alkaline, the 207nm peak of collagen red-shifted to 212nm indicating that the supra assembly of collagen misfolded with a tendency to forfeit its fibrillogenic structure. However, CH did not register a shift in its absorbance maxima, leading to the inference that the quasi-fibrillar structure of CH was retained even at pH 8. At pH 9 the amine peak red-shifted to 221nm for both CH and collagen depicting the loss of fibrillar property. This was due to the pH-induced negative charge gained by collagen leading to destabilization of the fibril.

The wavelength shifts of the amine group proved to be a favourable indicator of the hierarchical state of both CH and collagen. The results validated that amide N-H groups in both CH and collagen display a similar pH-induced absorbance shift confirming that peptides in CH were mimicking the folded hierarchy of the intact collagen. However, two marked differences were noticeable. At pH 4, CH existed in a less folded state than collagen, whereas at pH 8, CH managed to retain the fibrillar hierarchy even though collagen lost the same. This signifies that the peptides of CH subsisted in a more dynamic but stable quasi fibrillar state compared to that of its parent molecule, at least, in the alkaline pH region.

### 3.5 Secondary structure determination of CH

The circular dichroism spectroscopy measures the difference in absorption of right-handed and left-handed circular polarized light by the ensemble of peptide bonds constituting a protein backbone. Since the orientation of the bonds follows the secondary structure of the protein, CD spectroscopy becomes a powerful and accurate tool to determine the 3-D conformation of the protein [38,39]. Both CH and collagen displayed similar profiles as seen in Fig. 2c. At pH 7, CH and collagen exhibited negative peaks at 196-201nm indicating the presence of PP-II helix as the predominant secondary structure. Both samples displayed a negative value at 207nm and a shoulder like protrusion at 222nm. Changes in the 222nm wavelength have been reported to be linked with fibrillogenesis [40,41]. It has also been reported that negative signals between 204 and 210nm were a dominant indicator for the self-assembling tendency of a protein [40]. Based on the two parameters it was concluded that at pH 7, both samples exhibited a hierarchical fibrillar assembly. The data corroborated with the interpretations of the spectral shift analysis. It could be concluded that the peptides in CH wound around each other to form a mimic helix, and these mimic helices associateed through weak interactions to form a quasi-fibril.

### 3.6 Dimensions and ZP of CH

The dynamic Light Scattering (DLS) technique is based on the Brownian motion of dispersed particles. In DLS, an incident light beam scattered by a moving suspended particle is detected in a time-dependent fashion. This scattered light gives an estimate of the hydrodynamic radius of the particle [42].

As seen in Fig. 3a, CH displays a bimodal distribution indicating higher-order structure formation. The smaller distribution observed at 164.56±42.5nm particle size was due to an initial sidewise association of the mimic helices, that further assembled to form larger quasi-fibrils with average dimensions of 842.2±229nm. Collagen is known to display fibrillar properties at physiological pH, a property reflected in the bimodal size distributions of collagen. The smaller distribution was seen at 226.43±35.6nm matched the size of tropocollagen helices, while the second one was observed at 1260±1029nm, which accounted for the lateral associations of the tropocollagen units to form fibrillar structures.

**Fig. 3.**
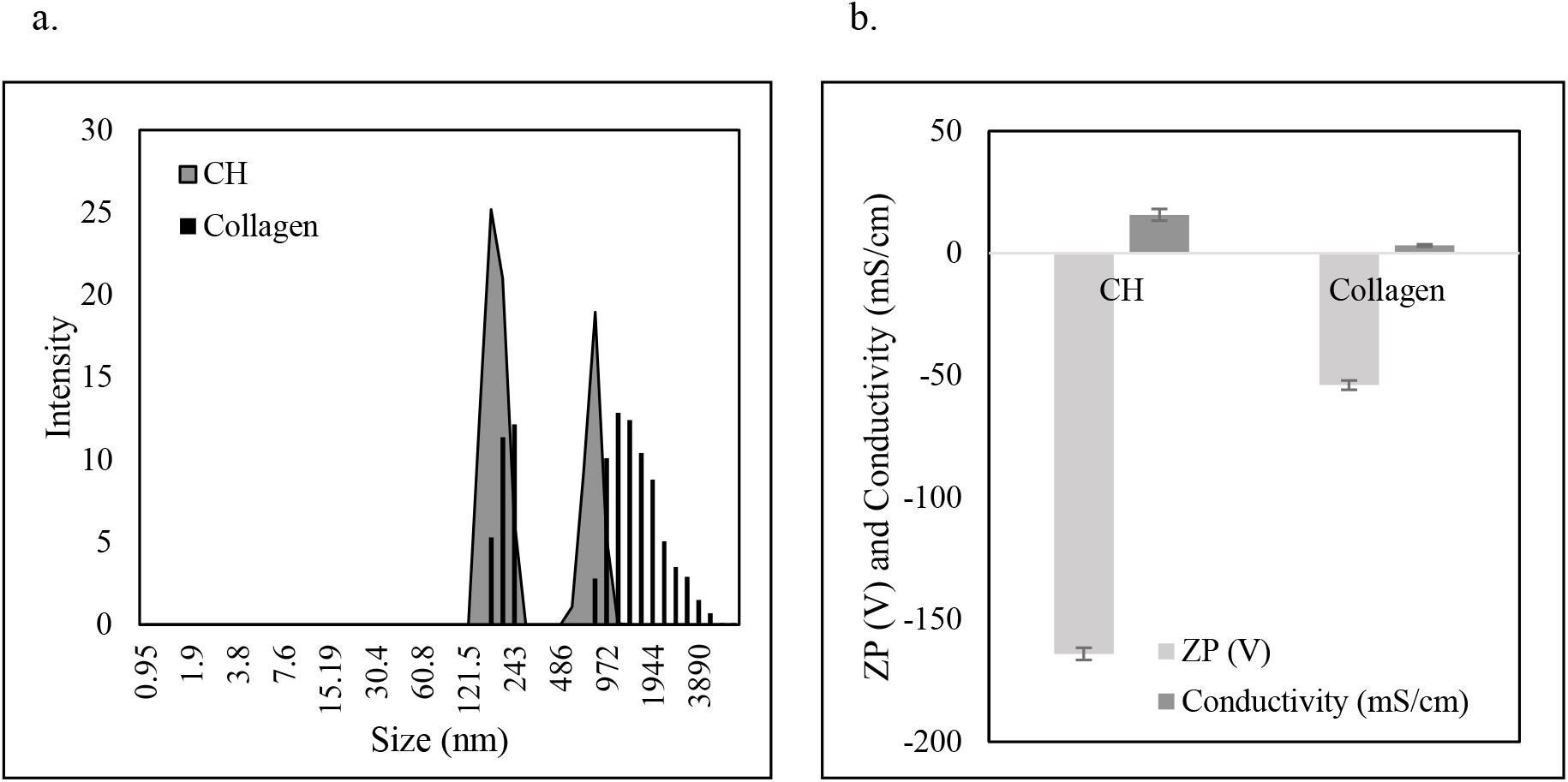
**a**- Distribution of particle diameter of CH (grey solid area) compared to collagen (lines). CH shows a small but definite presence of fibrillar assemblies **b** – CH displays a greater ZP and conductivity when compared to collagen

Zeta Potential (ZP) is a characterization technique employed to estimate the physical stability of a suspension by assessing the surface charge of molecules. A larger ZP value (positive or negative) is an indicator of better suspension stability [43]. The CH and collagen displayed ZP values of −164mV and −54mV respectively (Fig. 3b), indicating their propensity to remain in suspended form and not form aggregates. Overall, the DLS data complemented the data obtained from the spectral shift and CD studies.

### 3.7 Biomimetic growth of HA crystals on CH

The yield of HA from CH (CH/HA), based on the mean of multiple setups was determined to be 485.06±10.2mg. The control sample, HA grown on collagen (C/HA), yielded a lower amount of 468.02±14.5mg. This was of principal interest as the authors were unable to find any literature higher yield of HA crystals by using hydrolysate instead of protein or synthesized peptide. A thorough literature review also revealed that a lacuna existed regarding the comparison of HA nucleating capabilities of CH and collagen. Further structural characteristics were carried out to identify whether the CH could actually nucleate HA or the higher weight was merely due to the deposition of calcium phosphate.

### 3.8 Characteristics of CH/HA Nanocomposite

#### 3.8.1 Photoluminescence profile of CH/HA

Proteins, when excited by light with a wavelength ranging from 260nm to 600nm, exhibits fluorescence property [44]. Intrinsic fluorescence of marine collagen type-I is typically dominated by Phe, Tyr, and Trp residues at 282, 303, and 468nm respectively [45]. The luminescence profiles of the test composite, CH/HA, and the control, C/HA are given in Fig. 4a and 4b respectively. Both reflect the spectral signatures of the above-mentioned residues.

**Fig. 4.**
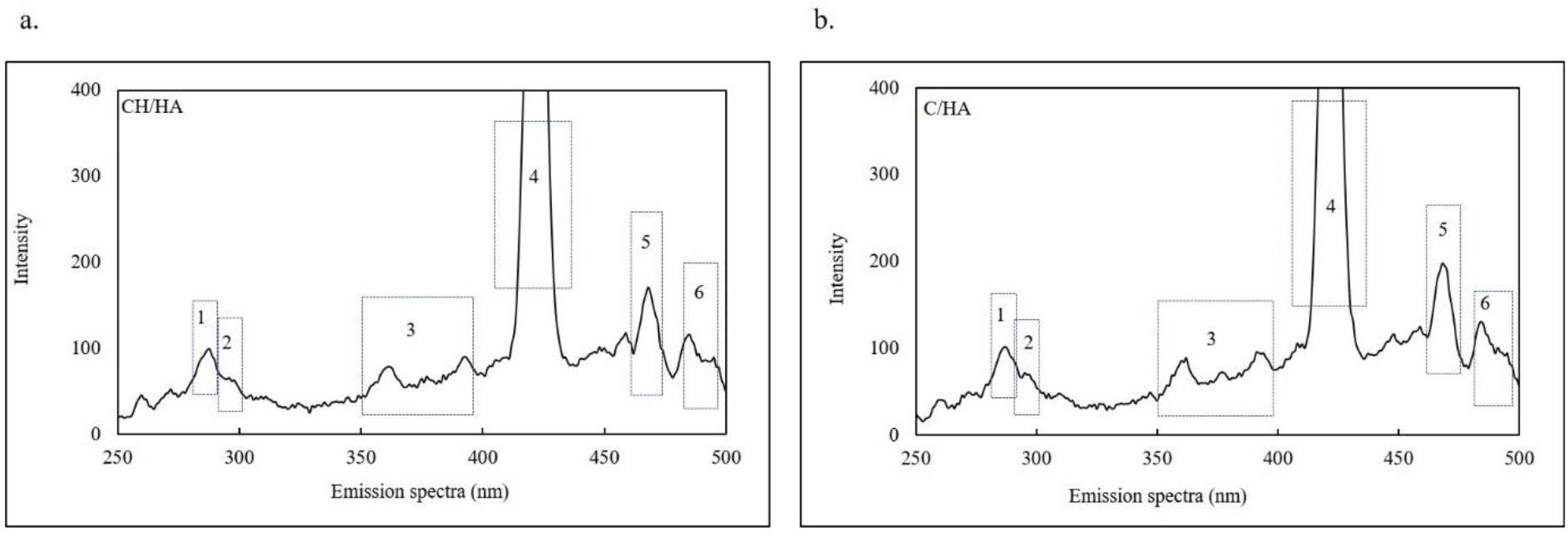
Photo Luminescence profile of a. CH/HA b. C/HA displaying peaks of specific amino acid residues and dityrosine crosslinks fon11ed due to HA nucleation. The highlighted peaks display the following amino acid residues (I) Phenylalanine (2) Tyrosine (3) Tyrosine aggregates (4) Dityrosine (5) Tryptophan coupled with histidine (6) Tryptophan

Phe and Tyr exhibited peaks at 284-285nm and 304nm respectively. Even though marine collagen has a higher abundance of Tyr, the low intensity observed in the luminescence profiles was due to collisional quenching with Trp residues located in close vicinity [44]. The peaks ranging from 307-393nm were attributed to the presence of Tyr aggregates, arising due to their close proximity in a hierarchical assembly [46,47]. The ubiquitous presence of closely spaced residues in both samples hinted at the presence of similar hierarchical fibrillar assemblies. The intense, sharp peak at 423-424nm depicted the presence of dityrosine cross bridges in both samples, a result of glutaraldehyde cross linking [48,49]. A prominent peak at 468nm in both spectra corresponded to the presence of a tryptophan aromatic ring coupled with the imidazole ring of a neighboring histidine [50]. Trp produced its distinct peak at around 490nm. Multiple low-intensity peaks at 443nm, 456nm, and 465nm in CH/HA corresponded to amino acid residue-mineral interactions.

Two distinct conclusions could be drawn from the close similarity of the CH/HA and C/HA peaks - (i) the peptides in CH were able to form a quasi-fibril that closely resembled collagen fibers (ii) CH was equally effective, if not better, at HA nucleation compared to collagen.

#### 3.8.2 IR Spectral profile of CH/HA

FTIR assists in structural investigations of specific functional groups by identifying the vibrations and the change in absorbance profile. Majority of these vibrations are localized on the amide and keto groups in the peptide bond. Their sensitivity to hydrogen bonding and dipolar interactions works as useful tools to identify changes in protein secondary structure. Fig. 5a and 5b represent the FTIR spectra of CH/HA and the control sample, C/HA.

**Fig 5.**
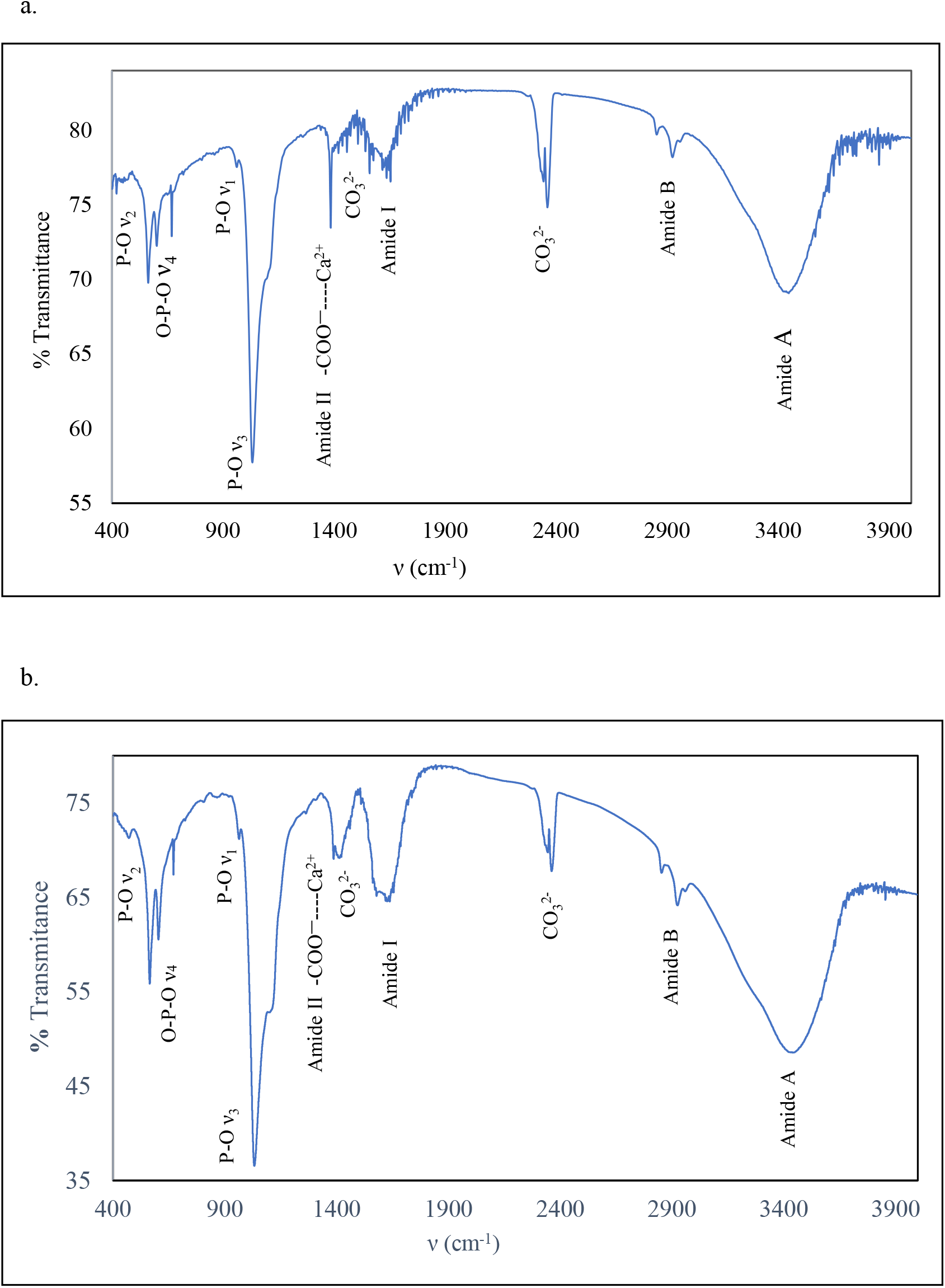
FTIR spectral profile of a. CH/HA b. C/HA exhibiting Amide A, B, I, II and HA-specific bands. CH/HA displays an intense peak at 1384cm^−1^ and lower Amide II indicating higher mineralization.

The Amide A bands arise due to the stretching vibration of O-H and N-H bond in the range of 3430-3900cm^−1^ [51,52]. CH/HA exhibited a broad Amide A in the range 3409-3462cm^−1^ while the C/HA exhibited a sharp high-intensity band at 3413.4cm^−1^. The Amide A peak provides substantial idea about the helical content of the substratum. An N-H stretch less than 3400cm^−1^ indicates that the peptide bonds are involved in intra-chain hydrogen bonding [53]. However, a shift to wave number greater than 3400cm^−1^, as seen for both CH/HA and C/HA was indicative of the lower abundance of hydrogen bonding, as expected for peptides existing in PP-II conformation. The Amide B peak representing >CH_2_ symmetric stretch at 2922cm^−1^ confirmed the presence of the protein substratum in both CH/HA and C/HA.

The Amide I band (1630-1680cm^−1^) occurs primarily due to the stretching vibration of the C=O group and it acts as a sensitive marker in the determination of the secondary structure of the protein [52]. The distortions of the C–N stretching and N–H bending vibrations help in identifying the environment of the bond and consequently, the secondary structure of the organic component. The peaks at 1641-1657cm^−1^ were prominent in both samples and indicated the presence of PP-II like coils. The band at 1665cm^−1^ was related to the hydrogen bonding propensity of hydroxyproline, a common component of collagen. The peak was significantly prominent in CH/HA in comparison to C/HA, owing to greater vibration in the more open quasi-fibrillar structure of the former in comparison to the tight packing of the latter. The presence of a sub-band near 1683cm^−1^signifies the degree of cross-linking, which is enormously imperative for mineralization [54]. The prominent visibility of 1684cm^−1^peak in CH/HA acts as evidence that the CH peptides were cross-linked more than that of collagen chains. A sub-peak at 1632cm^−1^ is known to indicate the presence of denatured coils [55]. CH/HA did not display this band indicating that the hierarchical structure was still maintained. However, CH/HA did show a band at 1638cm^−1^, which possibly indicated that it was less hierarchical than intact collagen.

The Amide II band occurs in the 1335-1402cm^−1^ and 1541-1558cm^−1^ positions. The former indicates the wagging motion of the proline ring –(CH_2_)- and the latter arises due to C-N stretching vibrations in combination with N-H bending [56]. Interestingly, during mineralization, the 1335cm^−1^ band is masked by a more prominent peak at 1383cm^−1^, occurring exclusively due to the interaction of Ca^2+^ with that of the carboxyl group of the protein/hydrolysate substratum. A prominent presence of this peak is considered as a prime marker for designating the state of mineralization [57]. The peak is observed in both nanocomposites, but is significantly larger in CH/HA, indicating a higher number of interactions between carboxyl groups and Ca^2+^ and consequently, better mineralization by the CH.

The second Amide II band (1541-1558cm^−1^) was different for both nanocomposites. CH/HA exhibited peaks at 1539cm^−1^ and 1547cm^−1^ which corresponded to the presence of triple helices whereas C/HA exhibited a single band at 1541cm^−1^, which indicated the presence of a distorted triple helix [58]. The Amide III peak has a band range of 1250-1350cm^−1^ and arises from the peptide bond N-H deformation coupled with N-C stretching. The band is barely visible at 1272cm^−1^ in CH/HA. A low amide III is indicative of a hierarchical fibrillar arrangement [59].

CH/HA and C/HA also exhibited typical peaks of phosphate. CH/HA displayed a sharp peak at 1033cm^−1^, indicating O-P-O (□_3_) bending [60]. It also exhibited a shoulder peak at 1104cm^−1^, a confirmatory peak for the presence of asymmetric HA. C/HA displayed a lower wavenumber band at 1029cm^−1^ and a shoulder peak at 1086cm^−1^. The 961-968cm^−1^ band corresponding to PO^−3^_4_ (□_1_) and the double band at 562cm^−1^ and 605cm^−1^ corresponding to PO^−3^_4_ (□_4_) were visible in both profiles. Both samples displayed a peak around 470cm^−1^ corresponding to PO^−3^_4_ (□_2_) stretch [60]. The presence of all four bands gave a strong indication that phosphate has been incorporated in the apatite formation.

The presence of carbonate peaks acts as indicators for determining the type of apatite formation. As seen in the figure, carbonate peaks were noticeable in the range of 1400-1500cm^−1^ and 2320-2370cm^−1^, designating the formation of type B apatite wherein a small number of phosphate ions are replaced by carbonate ions [17,61]. The increased carbonate ions might be from the atmospheric carbon dioxide which gets trapped into the lattice during crystal formation [62].

Earlier, the luminescence profiles confirmed that the CH existed as a higher-order fibrillar structure within CH/HA. The FTIR studies not only corroborated that but added further intriguing aspects. The CH/HA profile showed that the peptides existed as PP-II helices and even though presence of cross links was evident, the quasi-fibril formed by the CH peptides was somehow more open and dynamic, with a higher number of calcium-carboxyl interactions.

#### 3.8.3 XRD profile of CH/HA

XRD is a quantitative method to classify a crystal by detecting its characteristic planes of reflection. Fig. 6a depicts the XRD profile of CH/HA with sharp intense bands of HA overlapped with the broad amorphous phase of CH. A similar pattern was depicted by the control sample C/HA, although, with peaks of lower intensity (Fig. 6b).

**Fig. 6.**
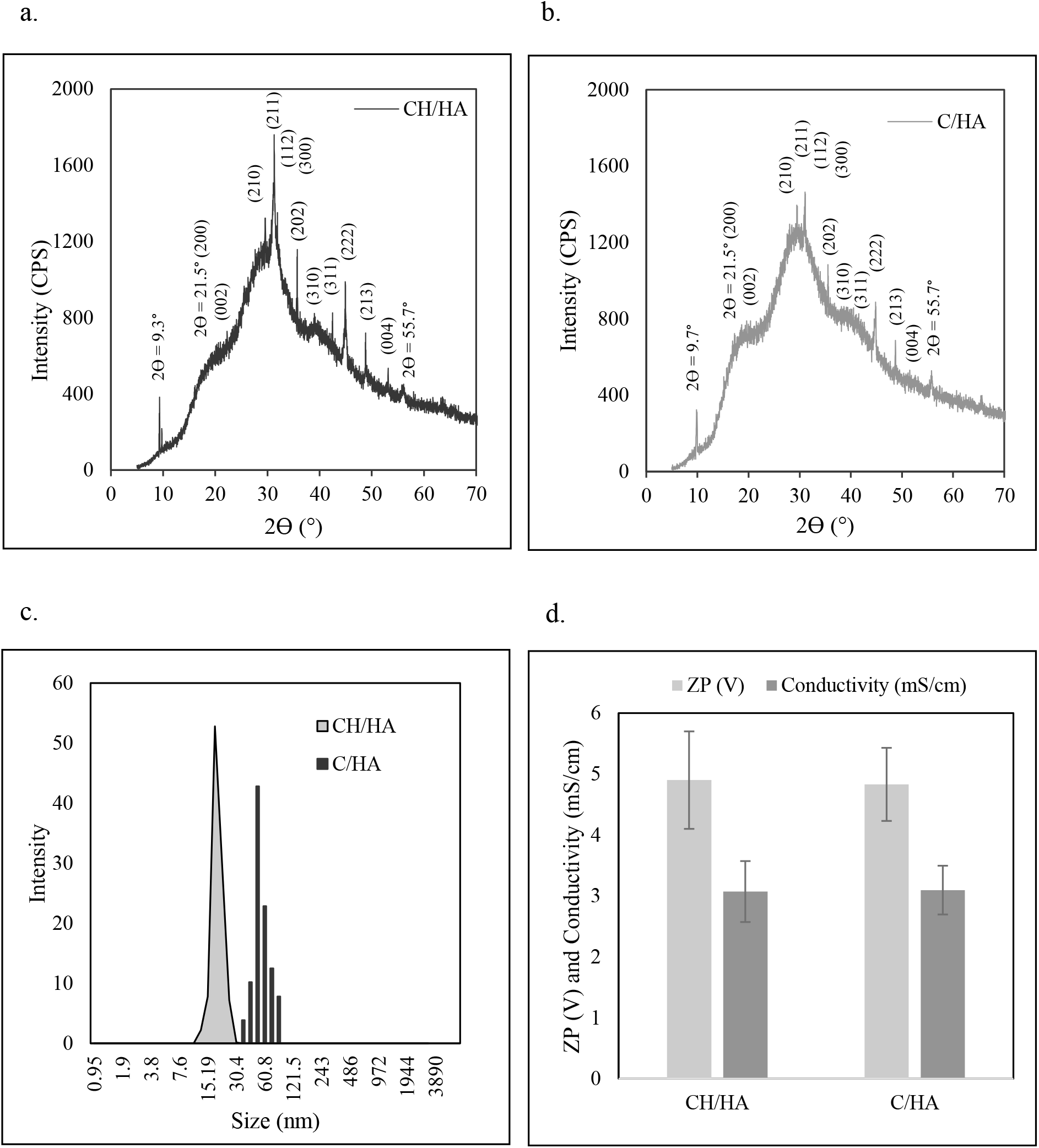
**a -** XRD spectral profile of CH/HA with greater crystallinity **b -** XRD spectra of C/HA **c -** Particle size distribution as observed by dynamic light scattering. CH/HA (grey, solid) displayed an average size of 41.16±13.21nm which was lower than the size distribution of C/HA (lines); 58.2±18.7nm. **d**- CH/HA and C/HA displayed a similar ZP (V) and Conductivity (mS/cm) value

The presence of high intensity peaks representing the planes 002 (2θ=25.9°), 210 (2θ =29.04°), 211 (2θ=31.8°), 112 (2θ=32.5°), 300 (2θ =32.9°), 202 (2θ=34.02°), 310 (2θ=39.8°), 311 (2θ=42.1°), 222 (2θ =46.7°), 213 (2θ =49.6°) and 004 (2θ =53.1°) coincided with HA planes listed in ICCD database confirming the presence of HA in CH/HA [63–65]. The control C/HA depicted similar planes with lower intensity indicating a lower crystalline state.

A sharp peak at 2θ=7° denotes the equatorial distance between PP-II helices coiled in a triple helical manner [56]. In this study however, both CH/HA and C/HA displayed a peak at 9.3 and 9.7° respectively, which corresponded to a *d* value of 0.93±0.028nm. It was evident that the formation of HA mineral phase was linked to an altered packing of the triple helices. The broad peak at 2θ = 21.57° (200) corresponds to a *d* value of 0.41nm. This peak is generally corroborated with amorphous scattering from multiple triple helices [66]. The peak was significantly diffused in CH/HA because the mimic-helical packing of CH was more dynamic and less sterically hindered than the intact triple helix of collagen. The larger peak at 2θ = 31.96° displayed a *d* value of 0.285nm which denoted the helical rise per residue in peptide chains oriented in a helical manner [67]. The peak at 55.7° represented a length of 0.17nm, which was attributed to the radius of a single PP-II helix. Similar peaks were observed in C/HA.

To conclude, CH displays an exquisite hierarchical assembly initiating with peptides oriented in PP-II conformation coiling with other PP-II peptides to form a triple-helical assembly. The smaller intermolecular packing distance along with a decidedly diffused 200 plane confirmed it to be a “mimic-helix” instead of an actual triple helix. These mimic-helices associated sideways to create a large quasi-fibrillar network, which becomes the substratum for HA nucleation and propagation. The sharper planes of CH/HA provided confirmatory evidence that crystallinity of CH/HA was better than collagen, confirming the superiority of CH quasi-fibril over collagen fibrils in HA nucleation.

#### 3.8.4 Size distribution of CH/HA

Fig. 6c describes the particle size distribution as measured by DLS. CH/HA displayed an average size of 41.16±13.21nm which was lower than the size distribution of C/HA; 58.2±18.7nm. The difference between the two samples was statistically significant (p<0.05, by t-test) indicating CH was able to nucleate crystals of smaller dimensions.

The ZP of CH/HA and C/HA ranged from +4.8 to +4.9mV (Fig. 6d). Biological macromolecules that exhibit ZP in the range of −25mV to +25mV are known to aggregate quickly in physiological fluids [43,68]. The composites exhibited low ZP indicating their low propensity to disassociate from a coating.

#### 3.8.5 SEM-EDX profile of CH/HA

The morphology of the CH/HA and C/HA nanocomposites are shown in Fig. 7a and 7b. The crystals exhibited a plate-like arrangement with serrated edges representing growing crystal planes [69,70]. Both nanocomposites depict a variety of size ranges. The CH/HA crystals recorded a small size range of 50-380nm while the C/HA crystals were larger (180-450nm). HA crystals with smaller dimensions effectively influence their biological and mechanical responses when used as coating materials [71,72]. This is one of the attractive features for the utilization of CH/HA as a coating material in comparison to C/HA.

**Fig 7.**
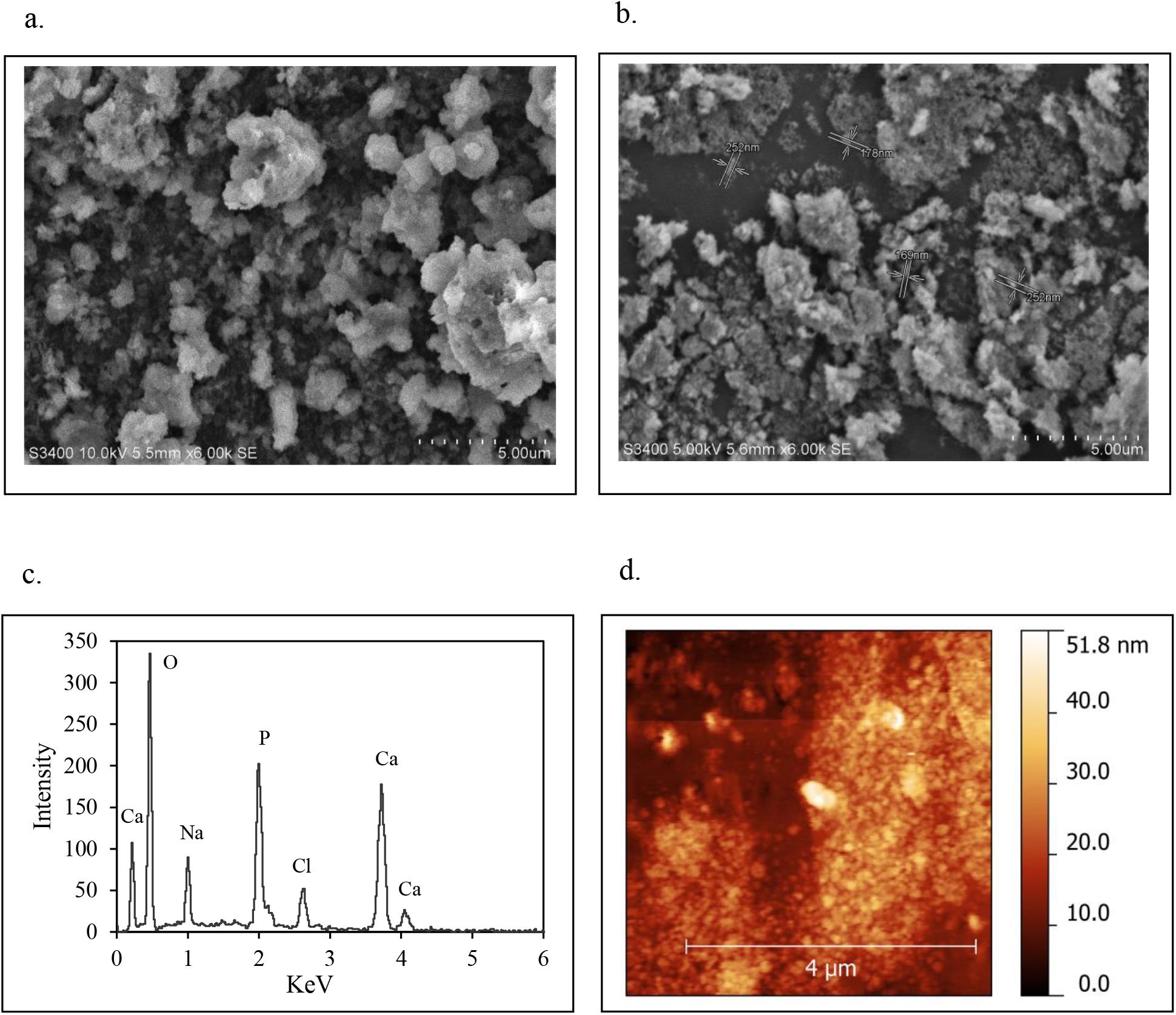
**a -** SEM image of coated CH/HA displaying smaller blooming crystals with the magnification of 1500X. The white dashed line represents a length of 5μm. **b -** SEM image of coated C/HA with the magnification of 1500X. The white dashed line represents a length of 5μm. **c**– EDX element analysis of CH/HA displaying a Ca/P ratio of 1.78 **d -** AFM analysis of CH/HA displaying the shape and nanophasic size of the HA crystals to be irregular oval-shaped with a height of 48±6.2nm. The white line represents a length of 4μm.

Ca and P exist in multiple phases such as amorphous calcium phosphates, brushite, monetite, tricalcium phosphate and hydroxyapatite. However, biomimetic synthesis ensures that the crystalline phase formed comprises majorly hydroxyapatite. The phases can be distinguished by the Ca/P ratio. Brushite and monetite exhibit a Ca/P ratio in the range of 1.1 to 1.3, whereas TCP and HA display a Ca/P ratio of 1.5 and 1.69 respectively [73]. The SEM-EDX data (Fig. 7c) reveals that the Ca/P ratio of CH/HA and C/HA nanocomposites in this study was 1.78 and 1.95 respectively. The Ca/P ratio of CH/HA is in close accordance with the 1.67 value for pure HA [74]. The Ca/P value of 1.78 for CH/HA matched with values obtained through previous biomimetic studies [17,75]. The observation of a small amount of Na in EDX results is in accordance with the published studies [76].

#### 3.8.6 Dimensional analysis of CH/HA

Accurate dimensions for the CH/HA crystals were obtained through AFM. As seen from Fig. 7d, the nanometric crystals exhibited an irregular oval shape with even smaller crystals scattered on the surface. The CH/HA crystals exhibited a size of 48±6.2nm, which matched the data obtained from DLS and SEM. Smaller particle sizes of constituent coating material enhance the surface roughness in the nanoscale. This property is important as pathogen-related contamination occurs less on nanosurfaces, giving the coating a distinctive advantage. As the surface roughness reduces from the sub-micron scale to the nano-scale level, two phenomena occur - the surface state gradually changes from hydrophobic to hydrophilic and becomes increasingly unfavorable for the adhesion of most bacteria. Second, the anchoring points for bacterial adhesion gradually decrease, weakening the bacterial-surface bonding strength [77,78]. At the same time, the nanometric roughness ensures integrin activation leading to higher osteoblast attachment [79]. Based on the size, it was expected that the CH/HA would work as a better osteoblast adhesion/ proliferation agent.

### 3.9 Osteoblast adhesion/ proliferation studies on CH/HA

The physicochemical properties of an implant surface, including hydrophobicity, surface free energy and roughness are an absolute necessity for selective osteoblast adhesion [77]. As seen in Fig. 8a, the amount coated had a significant effect (p<0.01) on the number of cells adhered. Overall, the adhesion exhibited a bell-shaped profile with maximum adhesion occurring around the 100-250μg range. This was expected from collagen-coated wells as excess collagen often eclipses the cell adhesion sites, reducing the number of cells that can adhere [80]. The nanocomposites followed a similar pattern. The cells adhered much better on the nanocomposites when compared to that of collagen-coated dishes, indicating the “osteophilicity” of the coat. A comparison between the adhesion capacity of CH/HA and C/HA displayed that overall, the former adheres to a higher number of cells. At 250μg CH/HA, the adhered cell count was significantly higher (p<0.05) than that of C/HA.

**Fig 8.**
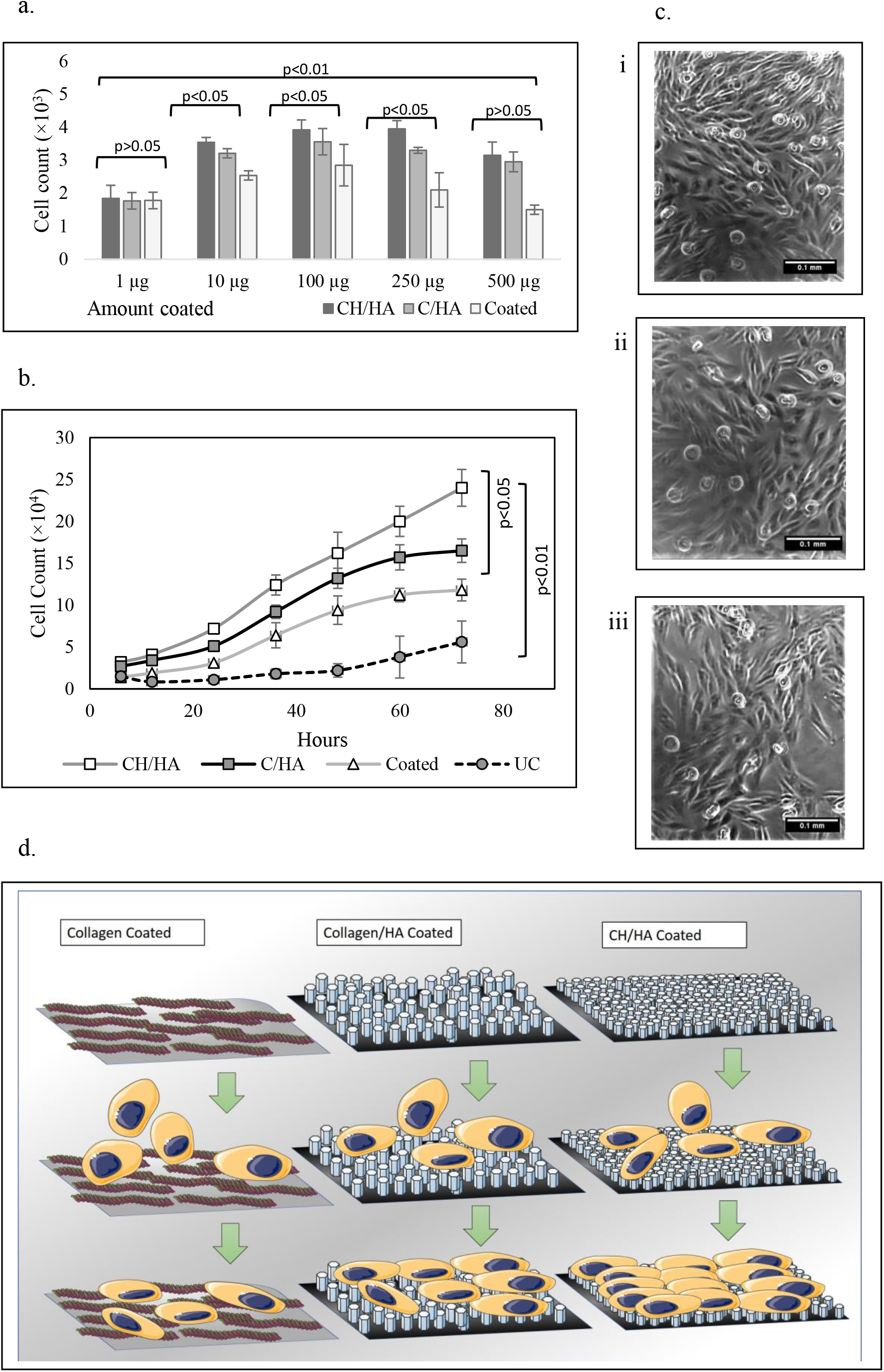
**a -** Osteoblast adhesion assay displaying a bell-shaped profile with the increasing amount of coating material. The significance values are displayed in the graph. **b -** Cell proliferation assay displaying a significantly higher cell growth when cultured in the presence of CH/HA when compared to C/HA. **c –** Photomicrographs representing MG63 growth on collagen-coated (I), CH/HA coated (II), C/HA coated (III) collagen coated wells. The white line represents a distance of 0.1mm. **d –** An illustration depicting the faster adhesion and greater proliferation of MG63 on the CH/HA coated surface when compared to C/HA and collagen-coated surfaces due to the smaller size of CH/HA making it more ‘osteophilic’.

Fig. 8b depicts the effect of nanocomposites on the rate of cell proliferation. Cells grown on nanocomposites and collagen displayed a normal growth rate after 20h. However, cells in the uncoated wells initiated a growth phase only after 36h. Cells proliferated significantly faster (p<0.01) on C/HA and CH/HA when compared to collagen-coated wells. Among the nanocomposites, cells growing on CH/HA were significantly higher (p<0.05) than C/HA. As seen in the figure, cells on CH/HA displayed exponential growth phase even at 72h. The proliferation rate on CH/HA was faster than C/HA by a factor of 1.39 and from coated dishes by a factor of 1.83. The doubling time of the cells on CH/HA and C/HA were 21.4 and 24.9 hours respectively while it took 28h for doubling on only collagen-coated wells. Fig. 8c provides the photomicrograph of cells grown till 60h with maximum cells observed on the CH/HA coated surface. Fig. 8d provides an illustration of the cell adhesion and proliferation occurring on three different coating surfaces. The osteoblast study corroborated previous reports that nanometric HA surfaces support higher osteoblast cell density [71,72] and provided evidence that HA grown on CH was a better osteophilic material than C/HA.

## 4. Conclusion

The study provided evidence that under *in vitro* conditions, collagen hydrolysate was able to nucleate nanophasic HA without the aid of any other externally added protein including collagen. The nanophasic CH/HA crystals exhibited higher yield and smaller dimensions when compared to the standard collagen-based nanocomposites and when used as a coating, was significantly better at osteoblast adhesion and proliferation.

The main reason for the high HA nucleation ability of CH was due to its intriguing quasi-fibrillar nature. The prevalence of proline constrained the peptides to orient in a PP-II conformation and coil with each other to form a triple-helical assembly. Owing to the smaller size (about 6 kDa), the triple-helical peptide fabrication was of diminished proportions with a reduced equatorial packing distance when compared to the collagen triple helix, and hence was coined as a “mimic-helix”. These mimic helices associated through hydrogen bonding to form a large quasi-fibrillar network that was more stable than collagen over the pH range of 6-8. At the same time, the CH fibrils were smaller in dimensions and more widespread in size, indicating their dynamic nature.

Structural studies on the nanocomposite of CH/HA revealed that the mimic-helix/ quasi-fibrillar hierarchy was maintained even after the hydrolysate was mineralized. The dynamic aspect of the CH quasi-fibril combined with a large number of C-terminals led to a massive number of exposed carboxylate groups that acted as calcium anchorage points more effectively than collagen, as evidenced by the presence of a distinctly prominent peak at 1383cm^−1^. At the same time, the high stability of quasi-fibrils assured that the groups maintained the orientation for a longer duration allowing crystallization to proceed. The addition of a minimum quantity of glutaraldehyde to the fibrils led to a large number of cross-links, as evidenced in the PLA and FTIR profiles, consequently contributing to the stability of the system.

A significant number of supportive markers demarcated the efficacy of the CH-based HA nucleation, the foremost of which was the higher yield when compared to collagen-based nucleation. The sharper HA-specific planes in CH/HA established the higher crystalline nature of CH/HA over that of C/HA. This was complemented by SEM and AFM results which yielded nanometric CH/HA crystals with smaller dimensions – an absolute necessity for osteoblast adhesion. The Ca/P ratio of CH/HA exhibited a value of 1.78 which was close to previous biomimetic HA and the value of pure HA (1.69). And to sum it all, the CH/HA nanocomposite was significantly better than collagen/HA in osteoblast adhesion and proliferation.

The present study furnishes the fact that the peptide-based dynamic quasi-fibril can act as better nucleating agents than the parent protein. This stand-alone property of CH demonstrates the absolute significance of protein secondary structure in biomineralization and provides an intriguing and eloquent insight into the biochemistry of collagen hydrolysate. Simultaneously, the results of this study provide substantial evidence for the “structural” theory, indicating that in vivo, the hierarchical assembly of an organic substratum, be it intact or hydrolzed, can both initiate and propagate the nucleation. The detailed morphological and analytical studies of nanocomposite act as a step forward in deciphering the qualitative and quantitative magnitude of this uniquely conserved protein in the field of biomimetics. The data acquired can act as a tie-in study to pursue the effectiveness of CH-based coatings on implants as an alternative to collagen-based coatings as a faster osseointegrative material.

## Acknowledgment

Some of the components of the illustrations in the graphical abstract and Fig. 8d have been taken from Servier Medical Smartarts. The authors thank the Chemistry Department of Dayananda Sagar University for their help in obtaining the FTIR profiles. The authors are grateful to Ms Prakruti Acharya for the immense help provided.

